# GPRuler: metabolic Gene-Protein-Reaction rules automatic reconstruction

**DOI:** 10.1101/2021.02.28.433152

**Authors:** Marzia Di Filippo, Chiara Damiani, Dario Pescini

## Abstract

**Background:** Metabolic network models are increasingly being used in health care and industry. As a consequence, many tools have been released to automate their reconstruction process *de novo*. In order to enable gene deletion simulations and integration of gene expression data, these networks must include gene-protein-reaction (GPR) rules, which describe with a Boolean logic relationships between the gene products (e.g., enzyme isoforms or subunits) associated with the catalysis of a given reaction. Nevertheless, the reconstruction of GPRs still remains a largely manual and time consuming process. Aiming at fully automating the reconstruction process of GPRs for any organism, we propose the open-source python-based framework `GPRuler`.

**Results:** By mining text and data from 9 different biological databases, `GPRuler` can reconstruct GPRs starting either from just the name of the target organism or from an existing metabolic model. The performance of the developed tool is evaluated at small-scale level for a manually curated metabolic model, and at genome-scale level for three metabolic models related to *Homo sapiens* and *Saccharomyces cerevisiae* organisms. By exploiting these models as benchmarks, the proposed tool shown its ability to reproduce the original GPR rules with a high level of accuracy. In all the tested scenarios, after a manual investigation of the mismatches between the rules proposed by `GPRuler` and the original ones, the proposed approach revealed to be in many cases more accurate than the original models.

**Conclusions:** By complementing existing tools for metabolic network reconstruction with the possibility to reconstruct GPRs quickly and with a few resources, `GPRuler` paves the way to the study of context-specific metabolic networks, representing the active portion of the complete network in given conditions, for organisms of industrial or biomedical interest that have not been characterized metabolically yet.

## Background

Current advances in genome sequencing technologies enable a fast and cheap overview into the genetic composition of virtually any organism. Nevertheless, determining the global metabolic profile of a cell or organism is fundamental to provide a comprehensive readout of its functional state, resulting from the interplay between genome, biochemistry and environment. In this context, genome-scale metabolic models (GEMs) offer a systemic overview for the investigation of cell metabolic potential, because of their key feature of embracing all available knowledge about the biochemical transformations taking place in a given cell or organism [1].

Over the years, extensive efforts have been made to automate the reconstruction process of these models, reaching high levels of quality and detail. However, less attention has been dedicated to clarify the associations among the set of genes involved in the catalysis of a given reaction.

According to the underlying catalytic mechanism, metabolic reactions can be classified as non-enzymatic and enzymatic. In the first scenario, metabolic reactions occur either spontaneously or small molecule catalyzed, implying that no gene is necessary for their catalysis. In the second case, enzymatic reactions occur only catalyzed by specific enzymes, which are protein macromolecules that are specifically responsible for reactions catalysis [2]. From the structural point of view, an enzyme may be classified as either monomeric or oligomeric entity. In its monomeric state, an enzyme consists of a single subunit, implying that a single gene is responsible of its final state. In this scenario, different protein isoforms representing highly related gene products differing in either their biological activity, regulatory properties, intracellular location, or spatio-temporal expression may alternatively catalyze the same function and, hence, the same reaction [3]. Conversely, in its oligomeric state, enzymes are protein complexes including multiple subunits that are all necessarily required to allow the corresponding reaction to be catalyzed.

In GEMs, the associations between genes, proteins and reactions and the description of how gene products concur to catalyze the associated reaction are usually encoded through logical expressions typically referred to as gene-protein-reaction (GPR) rules (Figure 1). GPR rules use the `AND` operator to join genes encoding for different subunits of the same enzyme, and the `OR` operator to join genes encoding for distinct protein isoforms of the same enzyme or subunit. Combining these two operators into expressions allow complex scenarios to be described, such as multiple oligomeric enzymes behaving as isoforms due to the sharing of a common part and to the presence of one or more subunits constituting distinctive features of the different isoforms.

**Figure 1.**
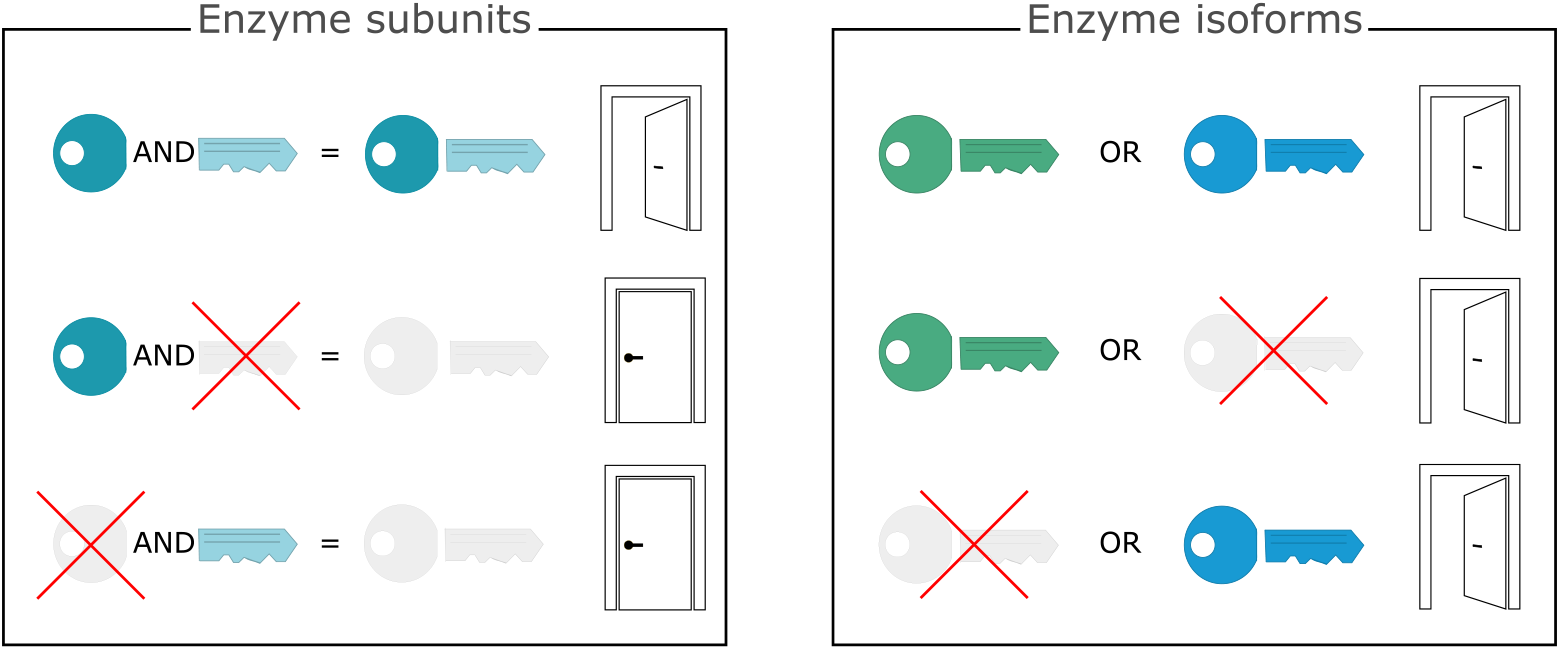
Logic of GPR rules. A metaphorical representation is exploited to explain the meaning of the `AND` and `OR` operators used in the reconstruction of GPR rules. In the box on the left named “Enzyme subunits”, the `AND` operator joins genes encoding for different subunits of the same enzyme. Metaphorically, the head and tail of the key represent the subunits of the same enzyme. Both parts of the key are required to open the door, and the lack of one of the two parts precludes the opening of the door. Biologically speaking, when both the subunits forming the enzyme are available, the enzyme can catalyse the reaction where it is involved. In the box on the right named “Enzyme isoforms”, the `OR` operator joins genes encoding for different isoforms of the same enzyme. In this case, two distinct keys representing the enzyme isoforms can alternatively open the door. Differently from the previous situation, just one of the two isoforms is sufficient to catalyse the reaction.

The reconstruction of GPR rules for their integration within metabolic networks does not constitute a novel task in literature. A Pubmed search of the last twenty years using the following four queries “`gene-protein-reaction rules`”, “`gene-protein-reaction associations`”, “`GPR rules`” and “`GPR associations`” returned 52 papers dealing with this issue. We reconstructed a network to analyse their citation status highlighting the most adopted strategies and data sources, as reported in Figure 2. Apparently, the most exploited sources are biological databases such as KEGG [4], UniProt [5], STRING [6] and MetaCyc [7], as well as genome annotations [8, 9], biochemical evidence presented in journal publications and reviews [10–12], and GPRs of closely related organisms [13]. The analysed articles also include work that relies exclusively on manual reconstruction of GPR rules, as e.g. [14], and work that limits to reconstruct one-to-one associations between genes and reactions rather than logical relationships among genes, proteins, and reactions [15].

**Figure 2.**
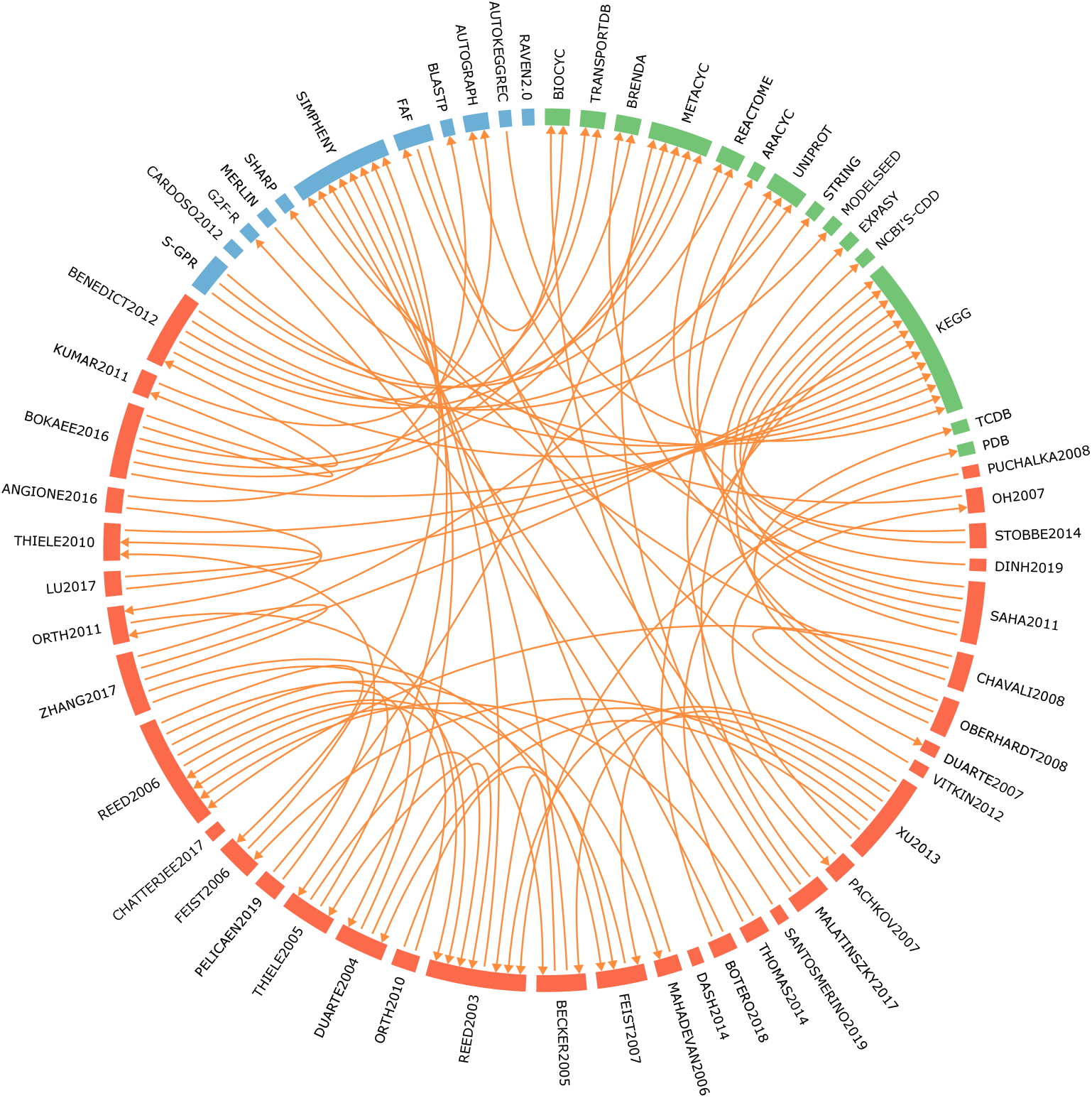
Literature analysis dealing with GPR reconstruction. In the circular plot, nodes correspond to the examined papers (red nodes), the adopted strategies (blue nodes) and data sources (green nodes). Directed edges connect a given work or tool to the exploited sources. If the source is not mentioned, the node remains isolated. Rectangles identify through their size the citation status of the corresponding node. References associated to each label are reported in the Additional file 1.

Our analysis of literature also highlighted the existing formally defined pipelines for the automatic generation of GPR rules within metabolic networks. The most cited one is *SimPheny* [16], which is a closed source industrial software application developed by the Genomatica company for the creation, enrichment and analysis of genome-scale metabolic models. In spite of its large use in literature, *SimPheny* is of difficult applicability, given that it is not released as independent product by the company. In view of its closed nature, it is not even possible to assess the strategy and information sources used by the tool, nor the automation level.

Two more recently implemented software have been made available that aim at reconstructing genome-scale metabolic networks for any target organism with the possibility to enrich them by creating the corresponding GPR associations: the *RAVEN Toolbox 2.0* [17] and *merlin* [18].

The *RAVEN* (Reconstruction, Analysis and Visualization of Metabolic Networks) *Toolbox 2.0* is a MATLAB-based tool for constraint-based metabolic modelling [17] that assists de novo semi-automated draft model reconstructions for given target organisms starting from genome sequence, including the possibility to automatically reconstruct their GPR rules. The complete pipeline can be executed starting from the genome sequencing of a given organism. Nevertheless, to our knowledge, the metabolic network draft returned by the tool does include information on the genes associated with each reactions but omits information on the relationships among the genes, which are joined together by `OR` boolean operators, regardless of whether their products are subunits or isoforms. Hence, substantial and time-consuming manual contribution is required to obtain the correct final GPRs.

The Metabolic Models Reconstruction Using Genome-Scale Information (*merlin*) is an open source software equipped with graphical interface to guide the user along all the proposed features in terms of genome-scale metabolic network reconstruction from genome annotation and implementation of the corresponding GPR associations through the usage of KEGG Orthology data. The graphical interface extends its applicability to a wide range of users since programming skills are not demanded. Nevertheless, during our testing, the software failed in automatically creating GPR associations for the target organism *Haloferax volcanii DS2* suggested in the tool tutorial, with empty rules associated to all tested reactions.

In view of the above, we propose a new methodology called `GPRuler` to efficiently automate the reconstruction process of GPR rules within metabolic networks of given organisms. In particular, we want to guarantee a white box among the currently available tools in order to introduce an open-source, clear and reproducible pipeline that can be applicable to any organism independently of its genome size, where the manual intervention is minimized in favour of an automatic information retrieval and managing. To achieve our aim, we rely on an extra available source not yet exploited by state of the art, which is the Complex Portal [19] database, which contain information about protein-protein interactions and protein macromolecular complexes established by given genes.

## Methods

### Tool implementation

`GPRuler` has been implemented in Python programming language, by using the 3.7 version. The overall `GPRuler` pipeline can be executed starting from two alternative inputs. In the first case, an already available draft SBML model or a simple list of reactions lacking the corresponding GPR rules can be provided. In the second case, `GPRuler` takes as exclusive input the name of the organism of interest. In both cases, the inputs are firstly processed to obtain the list of metabolic genes associated with each metabolic reaction in the target organism/model. This intermediate output is then used as input for the core pipeline, which returns as ultimate output the GPR rule of each metabolic reactions.

The overall proposed pipeline is graphically summarized in Figure 3, and its sequential execution steps are detailed in the following and in Figure 4.

**Figure 3.**
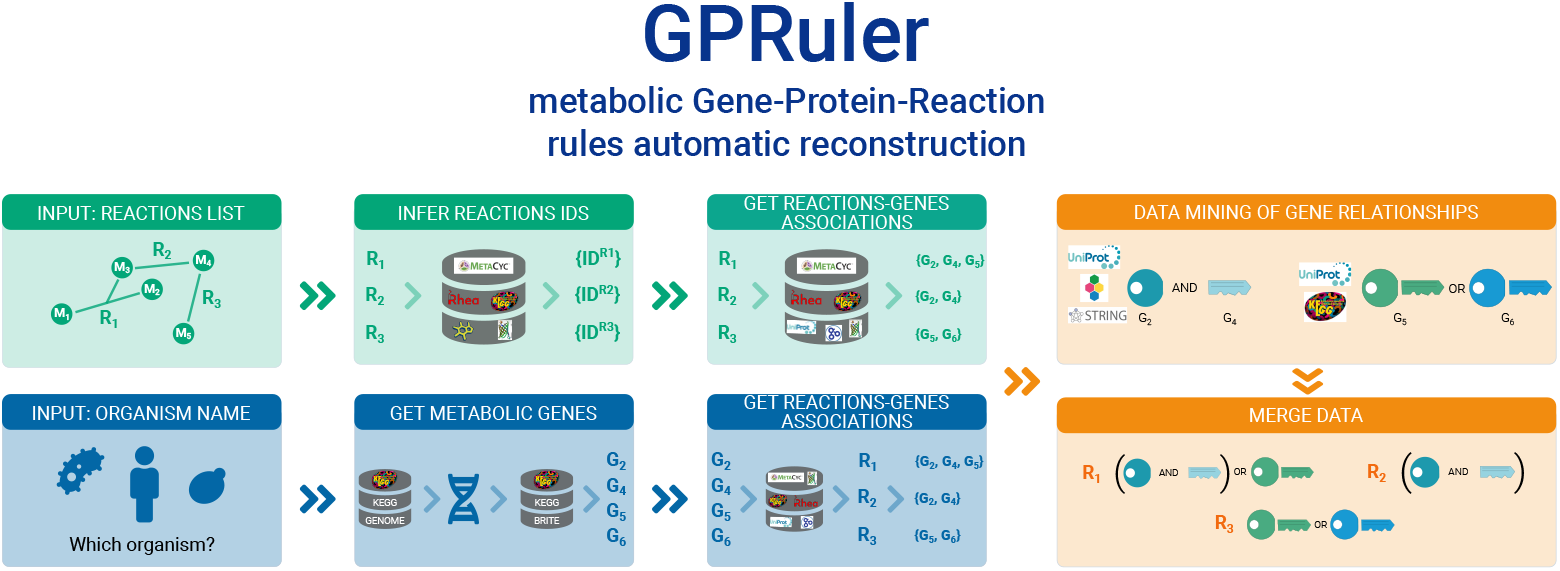
Graphical representation of `GPRuler` tool.

**Figure 4.**
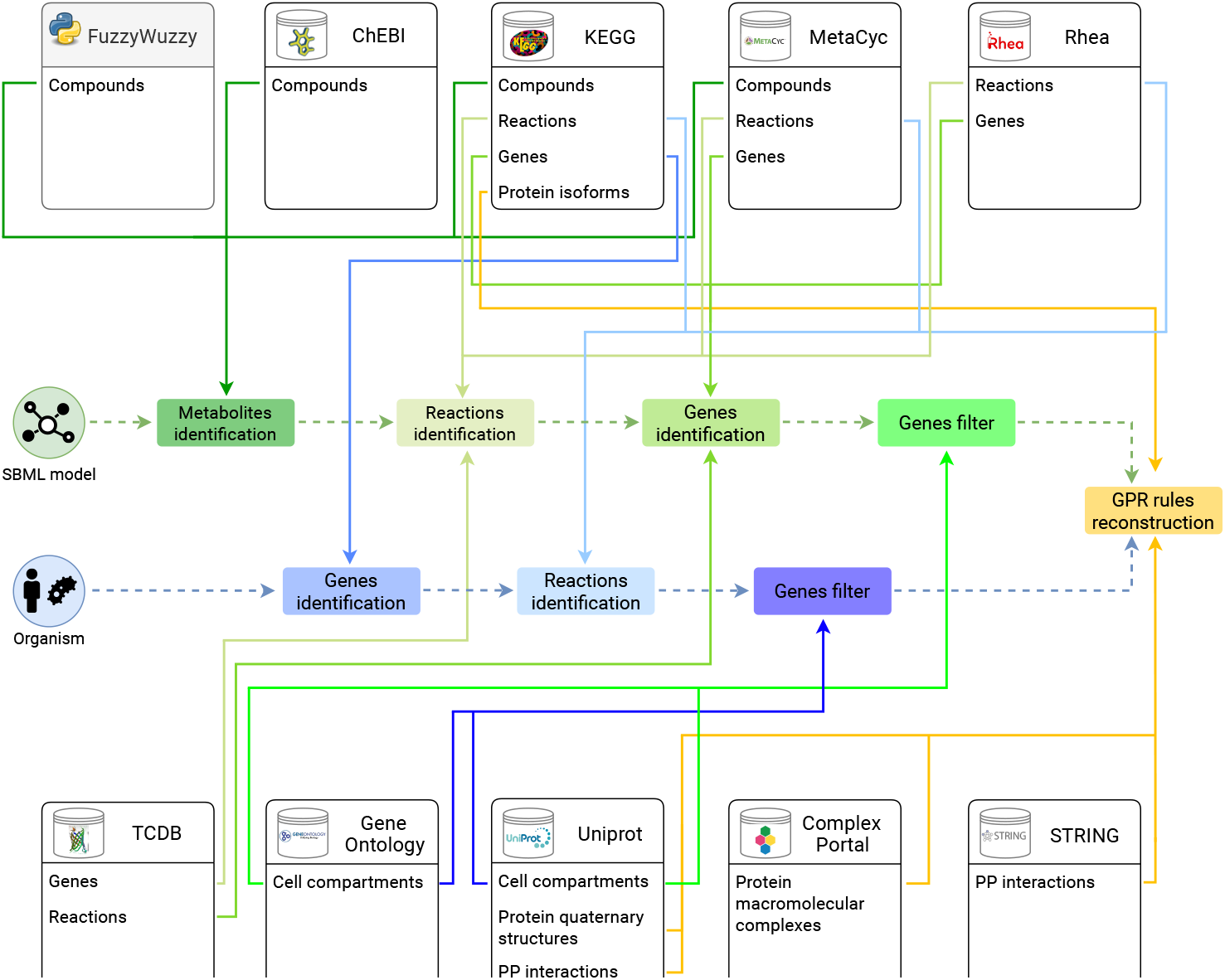
A detailed graphical representation of `GPRuler` tool. The central part of the figure illustrates the two alternative paths that can be followed to reconstruct the GPR rules according to the two possible inputs of `GPRuler`: the SBML model (green) and the organism name (blue). The green and the blue rectangles connected to each other by dashed arrows show the steps to follow in each path to achieve the core pipeline (orange rectangle), which returns as ultimate outcome the GPR rules. The ten boxes on the top and bottom of the figure represent the exploited sources used by `GPRuler`, including both biological databases (white boxes) and the FuzzyWuzzy Python package (gray box), listing which information is retrieved from each of them. Each coloured arrow links each step of the pipeline to the used source and, in particular, to the type of data for which that particular source is queried.

### From reactions list to annotated metabolic genes

`GPRuler` can handle the scenario where a metabolic model has already been reconstructed in terms of included biochemical reactions, but information on the associated genes and, of course, on GPR rules is not available yet.

In order to reconstruct this information, five biological databases are exploited: MetaCyc, KEGG, Rhea [20], the Chemical Entities of Biological Interest (ChEBI) [21], the Transporter Classification Database (TCDB) [22]. Firstly, all data available in these databases regarding the target organism, in their most recent version, are downloaded in order to speed up all the subsequent steps and, overall, the entire pipeline.

To get the set of genes involved in the catalysis of a given input reactions, cross-links identifiers to the above mentioned databases must be retrieved. To this aim, `GPRuler` identifies the metabolites involved in the reaction *in primis* (Figure 4, `Metabolites identification` green box), by using the ChEBI database as primary source, which is an open source dictionary about any distinguishable molecular entity naturally or synthetically produced involved in living organisms processes. Additionally to all the provided information about name, identifier, synonyms and direct links to the entries in KEGG and MetaCyc databases, we exploited two other information obtained from ChEBI: the IUPAC International Chemical Identifier (InChI) and the ChEBI ontology. The InChi is an alphanumeric string providing a unique digital signature to any compound based on its structural representation. The ChEBI ontology is a tree-like description of structurally related compounds, which are included in increasingly more general categories.

Metabolites are also identified in our pipeline through the MetaCyc database, which is a curated database of experimentally derived metabolic pathways involved in both primary and secondary metabolism, and of their associated metabolites, reactions, and genes. The PythonCyc package provided a Python interface to Pathway Tools, which is bioinformatics software accessing to all the information available in MetaCyc database.

Finally, KEGG COMPOUND and KEGG GLYCAN databases are queried to retrieve data about all the compounds annotated in this database, including links to external database such as ChEBI. KEGG is a biological resource consisting of multiple cross-linked databases that collect information relative to genes, enzymes, compounds, reactions and pathways.

Once the stored compounds data in ChEBI, MetaCyc and KEGG are extracted, the FuzzyWuzzy Python package is used for string matching between the target metabolite and a list of strings. Leveraging the Levenshtein distance to compute the similarity score between strings, the FuzzyWuzzy package extract the best matches to the target string.

Once the metabolites involved in each reaction of the input list have been assigned a proper identifier, `GPRuler` determines the reaction identifiers (Figure 4, `Reactions identification` green box), as annotated in KEGG REACTION, MetaCyc and Rhea database. Rhea is an expert-curated biochemical reactions resource using the chemical dictionary from ChEBI to describe reaction participants. Rhea is currently linked to several resources including UniProtKB, KEGG and MetaCyc.

In view of the fact that the three considered reaction databases are not completely independent one from another, as first action before querying them, our pipeline joins them in what we refer to as macro database. This macro database includes the union of the biochemical reactions present in each of the examined databases. In case the same reaction is found in multiple resources, it is represented as a unique object encompassing all the available annotated information about the reaction. Each reaction entry will include the corresponding set of reactants and products, cross-links to external resources and the list of corresponding EC numbers and catalyzing genes, from each of the consulted databases.

The macro database is then queried to infer for each input reaction the corresponding identifier and hence the corresponding annotated list of genes (Figure 4, `Genes identification` green box). Target reactions can be splitted in two kinds: internal, i.e. taking place within a given compartment inside the model, and transport, i.e. moving metabolites across compartments. For internal reactions, to retrieve the identifier it is sufficient to search for a reaction in the macro database that shares the same reactants and products.

For transport reactions, to retrieve the identifier it is necessary to query the TCDB database in addition to the macro database. The TCDB is a freely accessible database for transport proteins providing structural, functional, and evolutionary information about transporters from organisms of all types. This database is based on the Transporter Classification (TC) system, which is an approved classification system for membrane transport proteins analogous to the Enzyme Commission (EC) system used for the enzymes, except that it also includes functional and phylogenetic information. Exploiting the mapping provided by TCDB between the TC systems and the substrates annotated to be transported, TC codes for the target transport reaction are identified and the corresponding list of genes is isolated.

Finally, the identified list of genes of each reaction is then filtered to only keep entries whose subcellular location information is in accordance with those associated by the user to the reaction (Figure 4, `Genes filter` green box). Annotations about gene products localizations are inferred from Uniprot database and the Gene Ontology (GO) knowledgebase [23, 24], which provides an organism-independent description of genes and gene products in terms of biological processes to which the gene or gene product takes part, associated molecular functions and cellular components that refers to the cell location where a gene product is active. Each available annotation is associated to an evidence code indicating if the annotation is manually or computationally assigned to a particular term. Subcellular locations deriving from manual annotation are preferred over those deriving from automatic predictions.

### From organism name to the annotated metabolic genes and reactions

When the input is the name of a target organism, `GPRuler` seeks to retrieve all annotated metabolic genes of this organism, by accessing the KEGG database resource.

The first step of GPRruler is the determination of the KEGG identification code corresponding to the target organism’s name typed by the user. The user may be asked to disambiguate among a list of possible candidates codes.

The unambiguous identification of the target organism is then exploited to look for the corresponding list of genes involved in metabolic reactions catalysis (Figure 4, `Genes identification` blue box). In particular, the KEGG GENOME database is firstly queried to retrieve the complete reference genome sequences of the target organism. In order to isolate only metabolic actors, the KEGG BRITE functional hierarchy is used to filter out all the non-metabolic genes, whereas the KEGG REACTION database is queried to extract from the remaining ones the corresponding list of biochemical reactions involving them (Figure 4, `Reactions identification` blue box). Programmatical access to all data stored in KEGG database is performed thanks to the Bioservices Python package [25]. The macro database described in the previously section is also queried to enrich the selected list of reactions in terms of the corresponding annotated genes. Finally, the identified list of genes retrieved for each reaction is filtered according to their annotation of subcellular location from Uniprot and Gene Ontology, as already described (Figure 4, `Genes filter` blue box).

### Retrieving protein quaternary structures and protein-protein interactions from Uniprot

To characterize the relationships among the metabolic genes isolated in the previous steps of the pipeline and reconstruct the GPR rules (Figure 4, `GPR rules reconstruction` orange box), `GPRuler` relies on the information included in the Uniprot database and in particular in the “Interaction” section. Uniprot is an open-source database that provides a collection of manually and computationally determined functional annotations about proteins. Among the available data, the “Interaction” section organises in multiple subsections information about the quaternary structure of proteins and the set of binary protein-protein interactions established with other proteins or protein complexes. In particular, interchangeable participants and/or different chains or subunits belonging to the same complex are referenced through their gene names within a textual description.

For each metabolic gene associated with a reaction, `GPRuler` access the “Interaction” section of the Uniprot database to perform text mining, aiming at extracting, relatively to the input metabolic gene product, the list of genes participating in the same protein complexes, together with genes coding for isoforms. To guide the process of data extraction, we first identified the textual expressions containing relevant information. In this regard, we used the nltk Python library to tokenize the textual description of the “Interaction” section of investigated proteins and to carry out a frequency analysis, leaving aside the stop words that appeared frequently without conveying any meaning about the entire text. A frequency analysis of the remaining words highlighted the key words that, with the highest frequency count, guided in the extraction of the target information [26].

In particular, the analysis of a random set of genes belonging to two of the most annotated and known organisms, namely *Homo sapiens* and the yeast *Saccharomyces cerevisiae*, revealed the following four terms as top words: `complex`, `component`, `interact`, and `by similarity`. These key terms allow to identify, within the analysed texts, protein-coding genes that participate in the same protein complex of the queried protein, or that establish binary protein-protein interactions with it. These interactions are automatically derived from the IntAct Molecular Interaction Database [27], where all protein-protein interactions result from literature curation or manual user submission. In addition, manually curated information about protein-protein interactions derived from experimentally characterized proteins in closely related species and propagated to the target one, is also considered. In addition to the analysis of monograms, different length n-grams allowed to identify more extended expressions to enrich the search for target information in the “Interaction” section of Uniprot database. Among them, “`interact with`”, “`part of a complex with`”, “`consist of`”, “`associate with`”, “`heteromerization with`” represent some of the identified expressions.

In addition to the “Interaction” section, the “Function” section of the Uniprot database has been considered for text mining due to its role of describing isoform-specific functions or functionalities associated to the investigated protein as precursor of protein complexes. Information retrieval underwent the same initial text processing described ahead.

Programmatic access to data stored in Uniprot database is performed via the Bioservices package.

### Retrieving protein macromolecular complexes from Complex Portal

In addition to textual annotation of proteins, Uniprot database also offers cross-references to other biological databases storing data about protein-protein interactions.

Among them, Complex Portal is a manually curated database of macromolecular complexes relative to 21 key organisms representative of different taxonomic groups, including *Homo sapiens*, *Mus musculus*, *Saccharomyces cerevisiae*, *Escherichia coli*, *Arabidopsis thaliana*, *Caenorhabditis elegans*. All the included data derive from both physical and molecular interaction evidences resulting from literature search and manual inference from scientific background or homolog information in closely related species. Identifiers of protein complexes established by each input metabolic gene product are firstly extracted from the Uniprot search and access to Complex Portal data is achieved thanks to the Python requests and json libraries in order to extract all the components constituting the investigated complexes.

### Retrieving known and predicted protein-protein interactions from STRING

Another cross-referenced database to Uniprot about protein-protein interactions is STRING database. STRING is a database with a high coverage in terms of both annotated proteins and organisms, that deal with physical and functional associations for specific target proteins resulting from data integration of publicly available protein–protein interaction sources, including inference of functional associations between genes from genomic-context predictions, genome-scale laboratory experiments, gene co-expressions, co-citation analysis of scientific texts, manually curated interactions from other biological databases, knowledge transfer between organisms. The available knowledge is further complemented with computational predictions.

The programmatic access to STRING database is made possible by the STRING application programming interface (API). Among the implemented functionalities, “retrieving the interaction network” and “performing functional enrichment” are used to extract other protein-coding genes constituting a complex with the queried protein. In particular, the “retrieving the interaction network” method allows to retrieve the interaction network established by the target protein. Once this network is obtained, the “performing functional enrichment” method allows to perform functional enrichment analysis for each one of the selected proteins by mapping them onto several databases, including Gene Ontology, KEGG pathways, UniProt Keywords, PubMed publications, Pfam domains, InterPro domains and SMART domains. Thanks to this method it is possible to select, from the proteins included in the extracted interaction network, those belonging to the same protein complexes of the input queried protein.

### Retrieving protein isoforms from KEGG

Another type of relationship among genes involved in a given reaction that must be reconstructed is the possible existence of protein isoforms. In this regard, KEGG Orthology database, which belongs to the collection of databases included in KEGG database, is a database of functional orthologs, which have been defined based on their molecular functions by exploiting experimentally characterized genes or proteins with similar sequences in other organisms. From the KEGG gene identifier under investigation, it is possible to trace back to the associated KEGG Orthology identifier that returns for the specific organism under investigation the set of genes having this identifier, which are grouped together because of molecular functions similarity. Similarly to Uniprot, Bioservices package is used to programmatically access KEGG ORTHOLOGY database and automatically extract all the necessary information. In case that genes identifiers are not expressed as KEGG identifiers, a converter from external databases to KEGG database is provided by exploiting specific functionalities of Bioservices.

### Joining all retrieved information to generate final GPR rules

In the last step of `GPRuler`, all the data retrieved in the previous four steps is used to list, for each queried gene, the genes coding for subunits of the same protein complex to which the target gene belongs to or for alternative protein isoforms. Once this information is available, it needs to be organized to generate the final GPR rule. In this regard, all the possible pairs of genes associated to a given reaction are linked with either an `AND` or `OR` operator according to the type of relationship between them that has been previously inferred by `GPRuler`. In particular, the `AND` operator is assigned if the two genes encode for different subunits of the responsible enzyme, whereas the `OR` operator is assigned if they encode for alternative protein isoforms of the associated enzyme. In the simplest scenarios all the involved genes encode for isoforms of the same enzyme or they all encode for distinct subunits of the same enzyme. In a more heterogeneous scenario, in which both enzyme subunits and isoforms are present, GPRruler firstly joins genes related by an `AND` operator and enclose them in separated parentheses and then all the information about genes encoding for protein isoforms is included.

## Results

The core ability of `GPRuler` is to determine the GPR rule associated to each reaction. It follows that determining the right relationships among the involved genes (i.e., reconstruct the GPR rule) takes priority in validating the tool. To assess the degree of confidence of the reconstructed GPR rules and hence the accuracy of `GPRuler`, we exploited four curated metabolic models as ground truth.

According to the type of relationships established among genes involved in a given reaction, GPR rules can be categorized into five classes. In the simplest case, reactions can be associated to empty rules (here labelled as “No gene” rules) when no gene is involved in reaction catalysis, or to single gene rules (here labelled as “One gene” rules) when a unique gene is responsible for reaction catalysis. In these cases, minimum effort from `GPRuler` is required because the final GPR rule will simply correspond, respectively, to an empty string and to a rule involving the unique responsible gene.

An active role of our approach is played over more complex situations when multiple genes are involved in reaction catalysis generating the here labelled “Multi gene” rules, where the necessity to determine the right relationships among the involved genes takes place. In particular, “Multi gene” GPR rules can be characterized by just `OR` operators among genes (here labelled as “OR” rules), by just `AND` operators (here labelled as “AND” rules), or by both `AND` and `OR` operators in the most intricate scenarios (here labelled as “Mixed” rules).

We executed `GPRuler` by using a workstation equipped with 72 Intel Xeon 2.60GHz cpus and a TB of memory.

### Performance evaluation of GPRuler on a manually curated core model

We firstly assessed the accuracy of `GPRuler` in reconstructing the GPR rules of HMRcore model, which is a core model of central carbon metabolism that we extracted from the genome-wide HMR metabolic model [28] and we subsequently introduced and curated in [29–32]. We decided to use this model as benchmark to evaluate the performance of our approach because of the manually curated GPR rules associated to its reactions. Moreover, given the huge size of genome-scale models and the difficulty to control all the included GPRs, starting from a core model allowed a tighter control of the output.

HMRcore model consists of 315 reactions that, according to the previous GPR classification, are mainly associated to single gene rules representing the 45.7%. The remaining rules are 16.5% classified as “No gene” and 37.8% as “Multi gene”. The newly generated GPR rules has been compared with the corresponding ones in the original HMRcore model as follows. We reconstructed for each comparison the truth table undergoing the boolean expression of both the original and the new GPR rules by using the involved *n* genes as input variables. Considering that each variable can assume values equal to 1 (True) or 0 (False), the resulting truth table shows all the possible 2^*n*^ input combinations. Each input combination is then exploited to solve the two logic expressions under evaluation. After that, the resulting outputs are saved in a vector and add to the two truth matrices. The difference of the original and the newly generated GPR rule matrices will return 0 only when a perfect match between the two rules under comparison is obtained. In the opposite case, when a value different from 0 is obtained, a more or less wide discrepancy between the two input rules is achieved. It is worth noting that comparison of rules exceeding 20 genes needs to be manually evaluated because of the undergoing computationally expensive problem. The automated comparison is then followed by manual inspection of the negative matches in order to reveal false negatives due to cases where the rule generated by `GPRuler` updates the original counterpart in the tested model according to the biological knowledge from the explored databases.

In case of HMRcore model, the application of `GPRuler` on the reactions included in this model and the subsequent comparison with those stored in the original model, produced a perfect match for 110 of the 315 examined reactions, corresponding to a total of 34.9%. Considering the category of each rule, we correctly inferred 26.9% of “No gene” rules (14 out of a total of 52), 46.5% of the “One gene” rules (67 out of a total of 144), 15.4% of “AND” rules (2 out of a total of 13), 31.4% of the “OR” rules (27 out of a total of 86), and none of the 20 “Mixed” rules.

Following a manual deep analysis of the 205 wrongly obtained GPR rules (representing the 65.1% of total reactions of HMRcore model) compared to their original counterparts, 148 rules turned out to be replaceable with those generated through our methodology because in line with the information stored in the exploited databases. Thanks to this manual curation, the percentage of correctly predicted GPR rules of HMRcore increased from 34.9% to 81.9% (as shown in Figure 5A), reaching the 100% of the GPRs that can be automatically constructed. In particular, we curated 38 rules afferent to the “No gene” category by increasing the corresponding coverage to 100% (52 out of a total of 52), 68 rules afferent to the “One gene” category by increasing the corresponding coverage to 93.8% (135 out of a total of 144), 1 rule afferent to the “AND” category by increasing the corresponding coverage to 23.1% (3 out of a total of 13), 41 rules afferent to the “OR” category by increasing the corresponding coverage to 79.1% (68 out of a total of 86), and none of the 20 “Mixed” rules. In Figure 5B, a detailed overview of the automatically reconstructed rules is reported.

**Figure 5.**
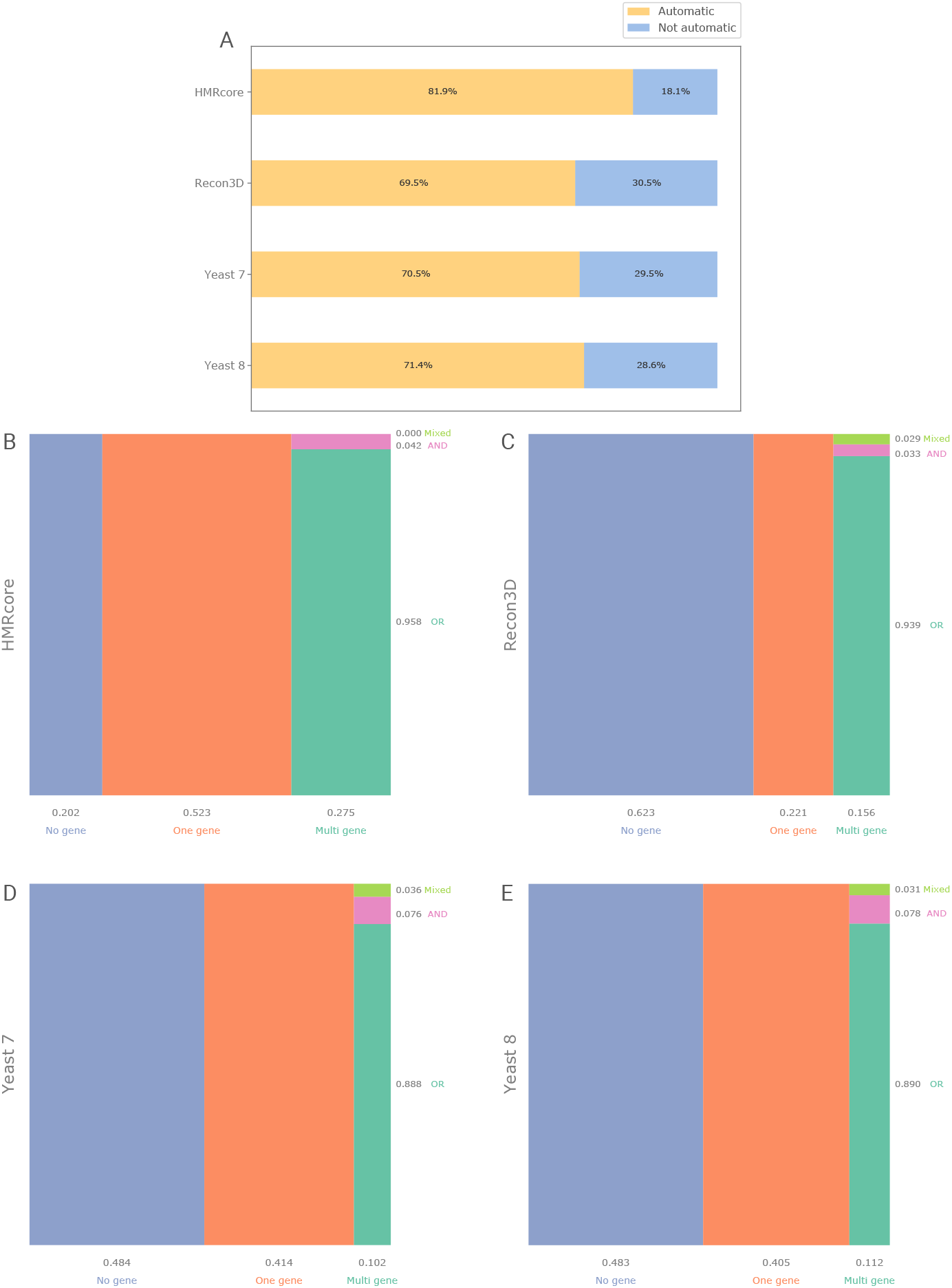
Performance of `GPRuler` tool. Panel A shows a summary of `GPRuler` performance highlighting the percentage of automatically reconstructed rules in each tested models (“Automatic” label coloured in dark yellow) against the not automatically reconstructed ones (“Not automatic” label coloured in light blue). The mosaic plots below show in B) HMRcore, C) Recon3D, D) Yeast 7 and E) Yeast 8 model the numerosity of the automatically reconstructed GPR rules as proportional to the size of internal rectangles, classified according to the type of relationships established among genes involved in reaction catalysis: “No gene”, “One gene” and “Multi gene”. In the “Multi gene” class, the three subclasses “OR”, “AND” and “Mixed” are also represented. On the horizontal side of each mosaic plot, the proportion of “No gene”, “One gene” and “Multi gene” automatically generated GPR rules in each model is reported. On the vertical side of each mosaic plot, the same information is reported for the three classes “OR”, “AND” and “Mixed” over the percentage of automatically generated Multi gene rules.

The execution time of GPRruler on HMRcore was 2 hours and 24 minutes. The complete list of GPR rules generated by `GPRuler` for HMRcore model is available in the Additional file 2.

### Performance evaluation of GPRuler on annotated genome-scale metabolic models

In light of the positive outcomes achieved with a small metabolic model of manually curated GPR rules, we extended the application of `GPRuler` at the genome-scale level for two of the most known and studied organisms, which are *Homo sapiens* and the yeast *Saccharomyces cerevisiae*. By considering the high level of knowledge of these organisms, we are confident in the accuracy and curation level of the GPR rules reconstructed for their metabolic reactions, whose generation derived from the integration of multiple biological databases, including SGD, BioCyc, Reactome, KEGG and UniProt.

The first considered genome-scale metabolic model is Recon3D [33], which is the most updated version of human metabolism. Recon3D model consists of 13543 reactions that, according to the previous GPR classification are mainly associated to single gene rules representing the 43.3%. The remaining rules consist of 28.2% “No gene” rules and 28.5% “Multi gene” rules.

The application of `GPRuler` on the reactions included in this model and the subsequent comparison with those stored in the original model produced a perfect match for 4378 of the 13543 examined reactions, corresponding to a total of 32.3%. In particular, considering the subcategory to which the obtained rules belong according to the included boolean operators, we correctly inferred 60.7% of “No gene” rules (3565 out of a total of 5867), 13.3% of the “One gene” rules (507 out of a total of 3821), 1.6% of “AND” rules (4 out of a total of 254), 8.9% of the “OR” rules (299 out of a total of 3345), and 1.2% of the “Mixed” rules (3 out of a total of 256).

Following the same procedure discussed in the previous section to understand the reason of the negative response returned by the remaining 9165 reactions (67.7%), we carried out a manual deep analysis by comparing the wrongly obtained GPR rules with their original counterparts. Unexpectedly, 5039 of the examined 9165 rules turned out to be replaceable in Recon3D model with those generated through our methodology because in line with the information stored in the exploited databases. Thanks to this manual curation, the percentage of total correctly predicted GPR rules of Recon3D increased from 32.3% to 69.5% (as shown in Figure 5A), reaching the 100% coverage of the GPRs that can be automatically reconstructed. More in detail, we curated 2301 rules afferent to the “No gene” category by increasing the corresponding coverage to 99.9% (5866 out of a total of 5867), 1574 rules afferent to the “One gene” category by increasing the corresponding coverage to 54.5% (2081 out of a total of 3821), 44 rules afferent to the “AND” category by increasing the corresponding coverage to 18.9% (48 out of a total of 254), 1081 rules afferent to the “OR” category by increasing the corresponding coverage to 41.2% (1380 out of a total of 3345), and 39 rules afferent to “Mixed” category by increasing the corresponding coverage to 16.4% (42 out of a total of 256). In Figure 5C, a detailed overview of the automatically reconstructed rules is reported.

A consideration about the examined rules in Recon3D model concerns some scenarios that should be revised in the original model independently of their comparison with rules generated from our approach.

A first case regards the two reactions labelled in Recon3D model as “`2OXOADOXm”` and “`AKGDm”`. GPR rules of these reactions did not result in completely accordance with the corresponding rule reconstructed through `GPRuler` due to the inclusion of the PDHX gene. A manual investigation of HumanCyc database [34] highlighted the affiliation of this gene to another protein complex different from the one involved in the reactions catalysis, which is the pyruvate dehydrogenase. Similarly, “`GLXO1”` reactions includes all genes corresponding to a different enzyme compared to the involved one. Consequently, inclusion of these genes in the current rules should be reviewed.

A second and consistent set of reactions was found to have contradictory GPR rules due to `AND` and `OR` operators simultaneously joining same genes. On behalf of this scenario, the reactions of Recon3D model labelled as “`AHEXASE2ly”`, “`AHEXASEly`”, “`NDPK10`”, “`NDPK1`” and “`PFK`” fall in this category. The most likely explanation of this behavior is the automatic generation of GPR rules successively missing a manual curation. Alongside with this hypothesis, a biological explanation of contradictory genes relationships might emerge in relation to alternative forms of the same enzyme. In these situations, the usage of parentheses within the logical expression becomes fundamental to discriminate automatic from biological annotations.

A last scenario emerged from the analysis of Recon3D model regards the reactions labelled as “`GUACYC`”, “`CREATt4_2_r`”, “`GLYKm`”, “`GUACYC`”, “`HPCLxm`”, “`NOS1`”, “`NOS2`”, “`r0400`”, “`r0924`”, “`MTHFC`”, “`PSP_L`”, “`TPI`”, “`r0345`”, “`GLYK`”, “`FTHFL`”, “`MTHFD2`”, “`HMR_2041`”, and “`HMR_4790`”. In this cases, comparison of the original GPR rule with `GPRuler` showed the need for a revision of the included gene identifiers. In all these reactions, genes no longer existing due to dismissal of their identifiers preventing the correct determination of the rule under consideration are included.

The execution time of `GPRuler` on Recon3D model was 231 hours and 33 minutes. The complete list of GPR rules generated by `GPRuler` for Recon3D model is available in the Additional file 3.

The other two genome-scale models considered for the performance testing of `GPRuler` are Yeast 7 and Yeast 8 models, which are the last two releases of *S. cerevisiae* genome-scale metabolism. The application of the pipeline to these two models returns comparable results between them.

Yeast 7 consists of 3493 reactions whose GPR rules resulted mostly categorized as “No gene” (34.1%) and “One gene” (48.7%). The remaining 17.2% consists of “Multi gene” rules. `GPRuler` applied to the reactions included in Yeast 7 proved to be able to perfectly reconstruct 1600 GPR rules, representing the 45.8% of the rules stored in the model. In particular, considering the category of each rule, we correctly inferred 69.6% of “No gene” rules (829 out of a total of 1191), 40.6% of the “One gene” rules (691 out of a total of 1701), 7.1% of “AND” rules (11 out of a total of 156), 17.1% of the “OR” rules (69 out of a total of 403), and none of the “Mixed” rules.

As in the previously discussed models analysis, manual investigation of generated GPR rules that negatively matched their counterpart in the original Yeast 7 model allowed to highlight all the cases where the newly reconstructed rule better fits the biology of the involved genes. Thanks to this manual curation, the global performance of `GPRuler` increased up to 70.5% (as shown in Figure 5A), totally covering the percentage of rule that can be automatically generated. The curation process of these 1893 rules revealed that 861 of them resulted better in line with the information stored in the exploited databases after their generation with `GPRuler`. More in detail, we curated 362 rules afferent to “No gene” category by increasing the corresponding coverage to 100% (1191 out of a total of 1191), 328 rules afferent to “One gene” category by increasing the corresponding coverage to 59.9% (1019 out of a total of 1701), 8 rules afferent to “AND” category by increasing the corresponding coverage to 12.2% (19 out of a total of 156), 154 rules afferent to “OR” category by increasing the corresponding coverage to 55.3% (223 out of a total of 403), and 9 rules afferent to “Mixed” category by increasing the corresponding coverage to 21.4% (9 out of a total of 42). In Figure 5D, a detailed overview of the automatically reconstructed rules is reported.

The execution time of `GPRuler` on Yeast 7 model was 14 hours and 2 minutes. The complete list of GPR rules generated by `GPRuler` for Yeast 7 model is available in the Additional file 4.

The updated model version of Yeast 7, namely Yeast 8, is a little bit larger than its predecessor since it consists of 3991 reactions. GPR rules classification showed that “No gene” and “One gene” rules cover, respectively, 34.5% and 47.4% of all the metabolic reactions included into the model, and “Mixed” rules cover the 18.1%. The execution of `GPRuler` perfectly reconstructed 1182 GPR rules, representing the 47.2% of the rules stored in the model. In particular, considering the category of each rule, we correctly inferred 74.9% of “No gene” rules (1032 out of a total of 1377), 39.7% of the “One gene” rules (751 out of a total of 1892), 5.5% of “AND” rules (9 out of a total of 163), 17.5% of the “OR” rules (90 out of a total of 513), and none of the “Mixed” rules.

The manual curation of the negatively matched GPR rules as in the previously presented models revealed, similarly to Yeast 7, the ability to correctly reconstruct the 71.4% of the model rules (as shown in Figure 5A), covering the 100% of rules that can automatically generated. Going into detail of the obtained GPRs, we correctly inferred 345 rules afferent to “No gene” category by increasing the corresponding coverage to 100% (1377 out of a total of 1377), 402 rules afferent to “One gene” category by increasing the corresponding coverage to 60.9% (1153 out of a total of 1892), 16 rules afferent to “AND” category by increasing the corresponding coverage to 15.3% (25 out of a total of 163), 194 rules afferent to “OR” category by increasing the corresponding coverage to 55.4% (284 out of a total of 513), and 10 rules afferent to “Mixed” category by increasing the corresponding coverage to 21.7% (10 out of a total of 46). In Figure 5E, a detailed overview of the automatically reconstructed rules is reported.

The execution time of `GPRuler` on Yeast 8 model was 13 hours and 30 minutes. The complete list of GPR rules generated by `GPRuler` for Yeast 8 model is available in the Additional file 5.

## Discussion

Regardless of the starting input used from `GPRuler` to reconstruct GPR rules, the accuracy of the generated output mainly depends on the level of annotation of the investigated organism within public biological databases.

Two main classes of errors emerged from the analysis of our results. The first class regarded the Uniprot database and the textual annotation of gene products it provides. This database groups any biological knowledge about a given gene product spanning from the general function to the annotated catalytic activity, to cross-references pointing to data collected from external databases. Moreover, a lot of the available information is hidden behind the annotated GO terms, which provide a textual description about biological processes and molecular functions associated to the gene product. All these data need to be explored to infer the correct biological function and biochemical reactions that gene products catalyse in order to solve the association between genes and reactions and how genes interact among them. However, analyses on text structures and on how information about protein function, structure and interactions is stored in the database revealed a lack of uniformity in the way data are stored and presented. This issue sometimes led to problems of failed mining of gene products annotations with the consequent lacking of the correct information within the generated rules. Most of `GPRuler` failures arose from the impossibility, unless by manual intervention, to codify upstream of the pipeline the associations between reactions and genes. Due to insufficient availability of data within the here exploited biological sources, `GPRuler` did not allow for an automated retrieval of these data, preventing from reconstructing the correct rule. In this context, Uniprot and Gene Ontology and the large quantity of information they contain could help to infer these relationships and generate the correct rules. Nevertheless, due to the lack of structured data in these databases, more work is required to process and understand the provided information. For this reason, sophisticated text mining algorithms will be evaluated for a more efficient data retrieval and interpretation. In addition, the success of `GPRuler` execution is also invalidated from the fact that much of the information we are looking for to reconstruct the right relationships are stored in organism-specific databases that we do not intend to include in `GPRuler` because the here presented pipeline has been developed to be applied to any organism. Consequently, only biological databases that are generic for all domains of life have been considered.

In the light of the above and as a consequence of the missing detailed description of the pipeline followed in the here tested metabolic models to generate the included GPR rules, there is reason to believe that most of the included data mainly come from manual curation.

In addition to all this, the pipeline has been conceived to be completely automatic, minimizing the manual contribution. When `GPRuler` starts from an existing metabolic model, how information is stored within models becomes crucial for the correct success of `GPRuler`. This specifically refers to the presence within the tested models of metabolites not immediately and automatically always traceable to the corresponding object into the considered biological databases, because of a missing standard in the used nomenclature that prevents from finding any link to existing compounds database. This issue pushed us to draw up a dedicated strategy in order to automatically identify metabolites involved in a given list of reactions, as described in Methods section. Nevertheless, due to the reasons explained above, this automatic procedure required in few specific cases a minimal manual contribution to review metabolites for which a shortlist of candidate identifiers is returned. In other situations, a manual curation of some particular instances has been instead necessary. An example is represented by the metabolite *ACP1* that corresponding to the Acyl-carrier protein should be included as *ACP* because it does not exist in the indicated form. Furthermore, we noticed that, within the tested models, reactions involving the NADPH cofactor are linked in some specific cases to reactions objects that within biological databases involve the adrenal ferredoxin metabolite. Following an analysis of biological knowledge, the adrenal ferredoxin results dependent from NADPH for its activity [35]. Therefore, we manually enriched the list of identifiers associated to NADPH metabolite with also those coming from adrenal ferredoxin. Similarly, the electron transferring-flavoproteins, which are FAD-containing proteins [36], have been manually linked to the FAD and FADH_2_ cofactors. Finally, the last two cases of manual curation regarded situations where the oxidation states of a given metabolite is missing resulting in an ambiguity of its identification, such as for the iron element that usually existing as +2 or +3 cations is used in Yeast 7 model to just indicate the ferrous ion form, and the usage in Yeast 7 and Yeast 8 models of the generic compound diglyceride for referring, as reported in KEGG COMPOUND database, to the specific chemical entity 1,2-Diacyl-sn-glycerol.

In addition to the just introduced scenarios, cases of impossibility to identify metabolites by any methods both automatic and manual occurred. Being the metabolites identification involved in the target reactions the starting point of `GPRuler`, an insufficiently curated metabolic model becomes limiting for the correct detection of the related catalysing genes and their relationships, affecting in this way the performance of the tool.

## Conclusion

In this work, we proposed an open-source tool called `GPRuler` in order to automate the reconstruction process of GPR rules for any living organism.

We applied the developed tool to four case studies, namely the HMRcore metabolic model and the genome-scale metabolic models Recon3D, Yeast 7 and Yeast 8, in order to assess the accuracy of `GPRuler` and the degree of confidence of the reconstructed GPRs.

By evaluating the resulting rules, as compared to their original counterparts, we verified the ability of `GPRuler` to reproduce the original GPR rules with a very high level of accuracy. After a manual curation of the comparisons producing negative matches between original and newly created rules, `GPRuler` proved to be able to reach a total coverage of the rules that can in principle be automatically reconstructed without any manual intervention.

The developed pipeline, besides producing good outcomes in terms of performance and accuracy, allowed to point out some issues in the original GPR rules stored in the investigated genome-scale models, especially in Recon3D, that need to be considered for future revisions of the models themselves.

With the advent of high-throughput technologies, an increasingly intensive generation of omics data occurred, paving the way towards the achievement of a system-level knowledge of cells. Although informative, relying only on these data is not enough to phenotypically characterize cells. In order to improve phenotypic predictions, it is appropriate to consider that a complete functional cell readout is not limited to gene expression analysis of cell. On the contrary, a characterization of its metabolic profiling, which represents the closest level of investigation to cell phenotype is required considering that a complex and non linear intracellular regulatory system occurs between gene expression and metabolic level.

Over years, several strategies have been proposed in order to integrate gene expression data into GEMs [31, 37–39] to derive context-specific networks representing the active portion of the complete network in given conditions. In this way, more biologically meaningful metabolic insights in distinct experimental conditions can be derived as a function of genes expression profiles encoding for subunits or isoforms of specific enzymes. Regardless of the approach developed to integrate omics data within GEMs, the success of these tools and the reliability of the formulated hypotheses strictly depends on the quality of the GPR rules included into the models. Hence, the importance of curating the GPR rules, whose good quality can also guarantee the successful outcome of multiple phenotype prediction experiments, including simulations of gene knock-out or gene expression variation, which allow to analyse the effect of these perturbations on the global system functioning.

Given the strong interest of life sciences research towards the omics data integration, having novel implements able to improve new knowledge discovery by boosting the performance of already existing computational tools involved at the forefront in this research line currently represents a priority in the scientific community. `GPRuler` may represent a breakthrough in this sense as complement of all those computational tools demanding, for their optimal functioning, a reliable implement able to reconstruct GPR rules for any organism of interest in an automated way and minimizing the manual intervention. Although, as highlighted in the Results and Discussion sections, some aspects need to be improved, today `GPRuler` represents a significant step beyond the state of art in the GPRs reconstruction field, paving the way towards future versions where a complete automation of the pipeline will take place.

## Supporting Information

### Additional file 1

List of references associated to each label in Figure 5.

### Additional file 2

#### GPR rules generated by GPRuler for HMRcore model

Each row shows in the column *Rxn* the reaction for which the GPR rule has been generated, in the column *rule_original* the corresponding original rule in the model, in the column *rule_GPRuler* the rule reconstructed by `GPRuler` and in the column *Evaluation* the output of the comparison between the rule in the *rule_original* and in the *rule_GPRuler* column: *Perfect match* if the comparison produced a perfect match between the two rules; *Corrected by GPRuler* if the two GPRs resulted different, but the original one in the model turned out to be replaceable by the rule generated by GPRruler because more in line with the underlying biology; *Not automatically reconstructed by GPRuler* if the GPR reconstructed by `GPRuler` did not result automatically attributable to the original form stored in the model.

### Additional file 3

#### GPR rules generated by GPRuler for Recon3D model

Each row shows in the column *Rxn* the reaction for which the GPR rule has been generated, in the column *rule original* the corresponding original rule in the model, in the column *rule_GPRuler* the rule reconstructed by `GPRuler` and in the column *Evaluation* the output of the comparison between the rule in the *rule original* and in the *rule_GPRuler* column: *Perfect match* if the comparison produced a perfect match between the two rules; *Corrected by GPRuler* if the two GPRs resulted different, but the original one in the model turned out to be replaceable by the rule generated by GPRruler because more in line with the underlying biology; *Not automatically reconstructed by GPRuler* if the GPR reconstructed by `GPRuler` did not result automatically attributable to the original form stored in the model.

### Additional file 4

#### GPR rules generated by GPRuler for Yeast 7 model

Each row shows in the column *Rxn* the reaction for which the GPR rule has been generated, in the column *rule original* the corresponding original rule in the model, in the column *rule_GPRuler* the rule reconstructed by `GPRuler` and in the column *Evaluation* the output of the comparison between the rule in the *rule original* and in the *rule_GPRuler* column: *Perfect match* if the comparison produced a perfect match between the two rules; *Corrected by GPRuler* if the two GPRs resulted different, but the original one in the model turned out to be replaceable by the rule generated by GPRruler because more in line with the underlying biology; *Not automatically reconstructed by GPRuler* if the GPR reconstructed by `GPRuler` did not result automatically attributable to the original form stored in the model.

### Additional file 5

#### GPR rules generated by GPRuler for Yeast 8 model

Each row shows in the column *Rxn* the reaction for which the GPR rule has been generated, in the column *rule original* the corresponding original rule in the model, in the column *rule_GPRuler* the rule reconstructed by `GPRuler` and in the column *Evaluation* the output of the comparison between the rule in the *rule original* and in the *rule_GPRuler* column: *Perfect match* if the comparison produced a perfect match between the two rules; *Corrected by GPRuler* if the two GPRs resulted different, but the original one in the model turned out to be replaceable by the rule generated by GPRruler because more in line with the underlying biology; *Not automatically reconstructed by GPRuler* if the GPR reconstructed by `GPRuler` did not result automatically attributable to the original form stored in the model.

## Availability of data and materials

`GPRuler` tool is available in the qLS GitHub repository at https://github.com/qLSLab/GPRuler. The datasets supporting the conclusions of this article are included within the article and its additional files.

## Acknowledgments

The institutional financial support to SYSBIO.ISBE.IT within the Italian Roadmap for ESFRI Research Infrastructures is gratefully acknowledged. Financial support from the Italian Ministry of University and Research (MIUR) through grant ‘Dipartimenti di Eccellenza 2017’ to University of Milano Bicocca, Department of Biotechnology and Biosciences is also greatly acknowledged.

## Author contributions

Conceptualization: MD and DP; Implementation of `GPRuler` tool: MD and DP; Manuscript writing: MD, DP and CD. All authors read and approved the final manuscript.

