## Additional File 1 for "GPRuler: metabolic Gene-Protein-Reaction rules automatic reconstruction"

### Table 1

This table reports the references associated to the labels present in figure 1.

| Label | Reference | DOI |
|---|---|---|
| PUCHALKA2008 | **Genome-scale reconstruction and analysis of the Pseudomonas putida KT2440 metabolic network facilitates applications in biotechnology.** PLoS computational biology, 4(10), 2008 | 10.1371/journal.pcbi.1000210 |
| OH2007 | **Genome-scale reconstruction of metabolic network in Bacillus subtilis based on high-throughput phenotyping and gene essentiality data.** Journal of Biological Chemistry, 282(39):28791--28799, 2007 | 10.1074/jbc.M703759200 |
| STOBBE2014 | Stobbe,M.D. et al. **Knowledge representation in metabolic pathway databases.** Briefings in bioinformatics, 15(3):455--470, 2014 | 10.1093/bib/bbs060 |
| DINH2019 | Dinh,H.V. et al. **A comprehensive genome-scale model for Rhodosporidium toruloides IFO0880 accounting for functional genomics and phenotypic data.** Metabolic engineering communications, 9:e00101, 2019. Elsevier. | 10.1016/j.mec.2019.e00101 |
| CHAVALI2008 | Chavali,A.K., et al. **Systems analysis of metabolism in the pathogenic trypanosomatid Leishmania major.** Molecular systems biology, 4(1), 2008. | 10.1038/msb.2008.15 |
| VITKIN2012 | Vitkin, E. and Shlomi, T. **MIRAGE: a functional genomics-based approach for metabolic network model reconstruction and its application to cyanobacteria networks.** Genome biology, 13(11):R111, 2012. | 10.1186/gb-2012-13-11-r111 |
| XU2013 | Xu,Z., Sun,X. et al. **Construction and analysis of the model of energy metabolism in E. coli.** PloS one, 8(1), 2013. Public Library of Science. | 10.1371/journal.pone.0055137 |
| PACHKOV2007 | Pachkov,M., et al. **Use of pathway analysis and genome context methods for functional genomics of Mycoplasma pneumoniae nucleotide metabolism.** Gene, 396(2):215--225, 2007. | 10.1016/j.gene.2007.02.033 |
| THOMAS2014 | Thomas,A., et al. **Network reconstruction of platelet metabolism identifies metabolic signature for aspirin resistance.** Scientific reports, 4:3925, 2014. | 10.1038/srep03925 |
| MAHADEVAN2006 | Mahadevan,R., et al. **Characterization of metabolism in the Fe(III)-reducing organism Geobacter sulfurreducens by constraint-based modeling.** Applied and Environmental Microbiology, 72(2):1558--68, 2006. | 10.1128/AEM.72.2.1558-1568.2006 |
| BECKER2005 | Becker,S.A. and Palsson,B.Ø. **Genome-scale reconstruction of the metabolic network in Staphylococcus aureus N315: an initial draft to the two-dimensional annotation.** BMC microbiology, 5(1):8, 2005. | 10.1186/1471-2180-5-8 |
| ORTH2010 | Orth,J.D. et al. **Reconstruction and use of microbial metabolic networks: the core Escherichia coli metabolic model as an educational guide.** EcoSal plus, 2005. | 10.1128/ecosalplus.10.2.1 |

| Label | Reference | DOI |
|---|---|---|
| THIELE2005 | Thiele,I. et al. **Expanded metabolic reconstruction of Helicobacter pylori (iIT341 GSM/GPR): an in silico genome-scale characterization of single-and double-deletion mutants.** Journal of bacteriology, 187(16):5818--5830, 2005. | 10.1128/JB.187.16.5818-5830.2005 |
| PELICAEN2019 | Pelicaen,R. et al. **Genome-scale metabolic reconstruction of Acetobacter pasteurianus 386B, a candidate functional starter culture for cocoa bean fermentation.** Frontiers in Microbiology, 10, 2019. | 0.3389/fmicb.2019.02801 |
| S-GPR | de Mas,I.M. et al. **Stoichiometric gene-to-reaction associations enhance model-driven analysis performance: Metabolic response to chronic exposure to Aldrin in prostate cancer.** BMC Genomics, 20(1):1--12, 2019. | 10.1186/s12864-019-5979-4 |
| CARDOSO2012 | Cardoso,J. et al. **An Algorithm to Assemble Gene-Protein-Reaction Associations for Genome-Scale Metabolic Model Reconstruction.** IAPR International Conference on Pattern Recognition in Bioinformatics, 118--128, 2012. | 0.1007/978-3-642-34123-6_11 |
| G2F-R | Osorio,D. et al. **Find and Fill Gaps in Metabolic Networks.** R package | https://CRAN.R-project.org/package=g2f |
| SHARP | Krishnakumar,S. et al. **SHARP: genome-scale identification of gene--protein--reaction associations in cyanobacteria.** Photosynthesis research, 118(1-2):181--190, 2013. | 10.1007/s11120-013-9910-6 |
| SIMPHENY | Price,N.D. et al. **Genome-scale microbial in silico models: the constraints-based approach.** Trends in biotechnology, 21(4):162--169, 2003. | 10.1016/S0167-7799(03)00030-1 |
| FAF | Green,M.L. and Karp,P.D. **Using genome-context data to identify specific types of functional associations in pathway/genome databases.** Bioinformatics, 23(13):i205--i211, 2007. | 10.1093/bioinformatics/btm213 |
| BLASTP | Altschul,S.F. et al. **Gapped BLAST and PSI-BLAST: a new generation of protein database search programs.** Nucleic acids research, 25(17):3389--3402, 1997. | 10.1093/nar/25.17.3389 |
| AUTOGRAPH | Notebaart,R.A. et al. **Accelerating the reconstruction of genome-scale metabolic networks.** BMC bioinformatics, 7(1):296, 2006. | 10.1186/1471-2105-7-296 |
| AUTOKEGGREC | Karlsen,E. et al. **Automated generation of genome-scale metabolic draft reconstructions based on KEGG.** BMC bioinformatics, 19(1):467, 2018. | 10.1186/s12859-018-2472-z |
| BIOCYC | Karp,P.D. et al. **The BioCyc collection of microbial genomes and metabolic pathways.** Briefings in bioinformatics, 20(4):1085--1093. 2019. | 10.1093/bib/bbx085 |
| TRANSPORTDB | Elbourne,L.D.H. et al. **TransportDB 2.0: a database for exploring membrane transporters in sequenced genomes from all domains of life.** Nucleic acids research, 45(D1):D320--D324. 2017. | 10.1093/nar/gkw1068 |
| BRENDA | Jeske,L. et al. **BRENDA in 2019: a European ELIXIR core data resource.** Nucleic acids research, 47(D1):D542--D549. 2019. | 10.1093/nar/gky1048 |
| REACTOME | Fabregat,A. et al. **The reactome pathway knowledgebase.** Nucleic acids research, 46(D1):D649--D655. 2018. | 10.1093/nar/gkv1351 |

| Label | Reference | DOI |
|---|---|---|
| ARACYC | Mueller,L.A. et al. **AraCyc: a biochemical pathway database for Arabidopsis.** Plant physiolog, 2(2):453--460. 2003. | 10.1104/pp.102.017236 |
| UNIPROT | UniProt Consortium. **UniProt: a worldwide hub of protein knowledge.** Nucleic acids research, 47(D1):D506--D515. 2019. | 10.1093/nar/gky1049 |
| MODELSEED | Henry,C.S. et al. **High-throughput generation, optimization and analysis of genome-scale metabolic models.** Nature biotechnology, 28(9):977. | 10.1038/nbt.1672 |
| EXPASY | Gasteiger,E. et al. **ExPASy: the proteomics server for in-depth protein knowledge and analysis.** Nucleic acids research, 31(13):3784--3788. 2003. | 10.1093/nar/gkg563 |
| NCBI'S-CDD | Marchler-Bauer,A. et al. **CDD: NCBI's conserved domain database.** Nucleic acids research, 43(D1):D222--D226. 2015. | 10.1093/nar/gku1221 |
| TCDB | Saier Jr,M.H. et al. **The transporter classification database (TCDB): recent advances.** Nucleic acids research, 44(D1):D372--D379. 2016. | 10.1093/nar/gkv1103 |
| PDB | Berman,H.M. et al. **The Protein Data Bank.** Nucleic acids research, 28(1):235-242. 2000. | 10.1093/nar/28.1.235 |
| REED2003 | Reed,J.L. et al. **An expanded genome-scale model of Escherichia coli K-12 (iJR904 GSM/GPR).** Genome biology, 4(9):R54, 2003. | 10.1186/gb-2003-4-9-r54 |
| REED2006 | Reed,J.L. et al. **Towards multidimensional genome annotation.** Nature Reviews Genetics, 7(2):130, 2006. | 10.1038/nrg1769 |
| FEIST2007 | Feist,A.M. et al. **A genome-scale metabolic reconstruction for Escherichia coli K-12 MG1655 that accounts for 1260 ORFs and thermodynamic information.** Molecular systems biology, 3(1), 2007. | 10.1038/msb4100155 |
| DUARTE2004 | Duarte,N.C. et al. **Reconstruction and validation of Saccharomyces cerevisiae iND750, a fully compartmentalized genome-scale metabolic model.** Genome research, 14(7):1298--1309, 2004. | 0.1101/gr.2250904 |
| FEIST2006 | Feist,A.M. et al. **Modeling methanogenesis with a genome-scale metabolic reconstruction of Methanosarcina barkeri.** Molecular systems biology, 2(1), 2006. | 10.1038/msb4100046 |
| ORTH2011 | Orth,J.D. et al. **A comprehensive genome-scale reconstruction of Escherichia coli metabolism—2011.** Molecular systems biology, 7(1), 2011. | 10.1038/msb.2011.65 |
| ANGIONE2016 | Angione,C. et al. **Multiplex methods provide effective integration of multi-omic data in genome-scale models.** BMC bioinformatics, 17(4):83, 2016. | 10.1186/s12859-016-0912-1 |
| BENEDICT2012 | Benedict, M.N. et al. **Genome-scale metabolic reconstruction and hypothesis testing in the methanogenic archaeon Methanosarcina acetivorans C2A.** Journal of bacteriology, 194(4):855--865, 2012. | 10.1128/JB.06040-11 |
| SANTOSMERINO2019 | Santos-Merino,M. et al. **New applications of synthetic biology tools for cyanobacterial metabolic engineering.** Frontiers in Bioengineering and Biotechnology, 7, 2019. | 10.3389/fbioe.2019.00033 |

| Label | Reference | DOI |
|---|---|---|
| DASH2014 | Dash,S. et al. **Capturing the response of Clostridium acetobutylicumto chemical stressors using a regulated genome-scale metabolic model.** Biotechnology for biofuels, 7(1):144, 2014. | 0.1186/s13068-014-0144-4 |
| DUARTE2007 | Duarte,N.C. et al. **Global reconstruction of the human metabolic network based on genomic and bibliomic data.** Proceedings of the National Academy of Sciences, 104(6):1777--1782, 2007. | 10.1073/pnas.0610772104 |
| MALATINSZKY2017 | Malatinszky,D. et al. **A comprehensively curated genome-scale two-cell model for the heterocystous cyanobacterium Anabaena sp. PCC 7120.** Plant physiology, 173(1):509--523, 2017. | 10.1104/pp.16.01487 |
| OBERHARDT2008 | Oberhardt,M.A. et al. **Genome-scale metabolic network analysis of the opportunistic pathogen Pseudomonas aeruginosa PAO1.** Journal of bacteriology, 190(8):2790--2803, 2008. | 10.1128/JB.01583-07 |
| ZHANG2017 | Zhang,Y. et al. **A new genome-scale metabolic model of Corynebacterium glutamicum and its application.** Biotechnology for biofuels, 10(1):169, 2017. | 10.1186/s13068-017-0856-3 |
| BOTERO2018 | Botero,K. et al. **A genome-scale metabolic model of potato late blight suggests a photosynthesis suppression mechanism.** BMC genomics, 19(8):863, 2018. | 10.1186/s12864-018-5192-x |
| LU2017 | Lu,H. et al. **Comprehensive reconstruction and in silico analysis of Aspergillus niger genome-scale metabolic network model that accounts for 1210 ORFs.** Biotechnology and bioengineering, 114(3):685--695, 2017. | 10.1002/bit.26195 |
| THIELE2010 | Thiele,I. and Palsson,B.Ø. **A protocol for generating a high-quality genome-scale metabolic reconstruction.** Nature protocols, 5(1):93, 2010. | 10.1038/nprot.2009.203 |
| BOKAEE2016 | Nazem-Bokaee,H. et al. **Assessing methanotrophy and carbon fixation for biofuel production by Methanosarcina acetivorans.** Microbial cell factories, 15(1):10, 2016. | 10.1186/s12934-015-0404-4 |
| SAHA2011 | Saha,R. et al. **Zea mays iRS1563: a comprehensive genome-scale metabolic reconstruction of maize metabolism.** PloS one, 6(7), 2011. | 10.1371/journal.pone.0021784 |
| KUMAR2011 | Kumar,V.S. et al. **Metabolic reconstruction of the archaeon methanogen Methanosarcina Acetivorans.** BMC systems biology, 5(1):28, 2011. | 0.1186/1752-0509-5-28 |
| CHATTERJEE2017 | Chatterjee,A. et al. **Reconstruction of Oryza sativa indica genome scale metabolic model and its responses to varying rubisco activity, light intensity, and enzymatic cost conditions.** Frontiers in plant science, 8:2060, 2017. | 10.3389/fpls.2017.02060 |
| MERLIN | Dias,O. et al. **Reconstructing genome-scale metabolic models with merlin.** Nucleic acids research, 43(8):3899--3910, 2015. | 10.1093/nar/gkv294 |
| RAVEN2.0 | Wang,H. et al. **RAVEN 2.0: A versatile toolbox for metabolic network reconstruction and a case study on Streptomyces coelicolor.** PLoS computational biology, 14(10):e1006541, 2018. | 10.1371/journal.pcbi.1006541 |
| KEGG | Kanehisa,M. et al. **KEGG as a reference resource for gene and protein annotation.** Nucleic Acids Research, 44(D1):D457--62, 2016. | 10.1093/nar/gkv1070 |

| Label | Reference | DOI |
| --- | --- | --- |
| METACYC | Caspi,R. et al. **The MetaCyc database of metabolic pathways and enzymes-a 2019 update.** Nucleic Acids Research, 48(D1):D445--D453, 2020. | 10.1093/nar/gkz862 |
| STRING | Szklarczyk,D. et al. **STRING v11: protein--protein association networks with increased coverage, supporting functional discovery in genome-wide experimental datasets.** Nucleic acids research, 47(D1):D607--D613, 2019. | 10.1093/nar/gky1131 |
